# Moonlight: a tool for biological interpretation and driver genes discovery

**DOI:** 10.1101/265322

**Authors:** Antonio Colaprico, Catharina Olsen, Claudia Cava, Thilde Terkelsen, Tiago C. Silva, Andrè Olsen, Laura Cantini, Gloria Bertoli, Andrei Zinovyev, Emmanuel Barillot, Isabella Castiglioni, Houtan Noushmehr, Elena Papaleo, Gianluca Bontempi

## Abstract

Cancer is a complex and heterogeneous disease. It is crucial to identify the key driver genes and their role in cancer mechanisms with attention to different cancer stages, types or subtypes. Cancer driver genes are elusive and their discovery is complicated by the fact that the same gene can play a diverse role in different contexts. Key biological processes, such as cell proliferation and cell death, have been linked to cancer progression. Thus, in principle, they can be exploited to classify the cancer genes and unveil their role. Here, we present a new method, Moonlight, that exploit expression data to classify cancer genes. Moonlight relies on the integration of functional enrichment analysis, gene regulatory networks and upstream regulator analysis from expression data to score the importance of biological cancer-related processes taking into account either the inter- or intra-tumor heterogeneity. We then employed these scores to predict if each gene acts as a tumor suppressor gene (TSG) or as an oncogene (OCG). Our methodology also allow to predict genes with dual role, i.e. the moonlight genes (TSG in one cancer type or stage and OCG in another), as well as to elucidate the underlying biological processes. Availability: https://bioconductor.org/packages/MoonlightR & https://github.com/ibsquare/MoonlightR/

## Introduction

Cancer development is influenced by mutations in two distinct categories of genes, known as tumor suppressor genes (TSG) and oncogenes (OCG). The occurrence of mutations in TSGs generally leads to increased cell proliferation, whereas mutations in OCGs could alter or increase their biological activity.

The gain-of-function of OCGs together with the loss-of-function of TSGs determine the processes that control the tumor formation and development (Hahn and Weinberg, 2002).

The Cancer Gene Census (CGC) (Futreal et al., 2004) is a catalog of somatic mutations that provides a list of currently known cancer driver genes following the OCGs / TSGs definition. The CGC contains some 600 genes implicated in cancer, but more recently, the availability of different “omics” technologies from large international consortia, such as the The Cancer Genome Atlas (TCGA) and Gene Expression Omnibus (GEO) have suggested many additional genes that may be involved in cancer and have brought new challenges to the field of cancer research (Vogelstein et al., 2013). These consortia included data of somatic mutations, gene expression, methylation and copy number alterations, along with clinical data. They share the ambitious goal of detecting crucial genes and molecular mechanisms underlying tumorigenesis in any major cancer type (Cancer Genome Atlas Research Network, 2008).

Several methods implemented so far used mutation data to detect candidates cancer drivers (Davoli et al., 2013), (Schroeder et al., 2014) (Schroeder et al., 2014; de Souza et al., 2014)) but although all these methods were demonstrated as effective, it remains critical to clarify the consequences of each mutation and to link these patterns with possible underlying biological interpretation. For example, some mutations bring premature stop codon and carry out reduced dosage of mRNA transcripts. Other mutations affect protein activities through can act as either an oncogene or tumor suppressor gene in different cancer types or during cancer progression.

For instance, most of the recent studies demonstrated that DEAD-box RNA helicase 3 (DDX3) acts as an oncogenic role in breast cancer (Bol et al., 2013), but some studies (Su et al., 2015) (Zhao et al., 2016) also showed that its downregulation promotes metastasis in colorectal cancer indicating a potential role as TSG.

Another example of a moonlight gene is represented by MMP12, a member of the matrix-metalloproteinase (MMP) family. The MMPs cause degradation of the extracellular matrix and basement membranes, contributing to the pathogenesis of tissue destructive processes in a wide variety of diseases. It has been proposed as an oncogene in breast cancer, (Hegedüs et al., 2008), (Shin et al., 2005), hepatocellular carcinoma (Ng et al., 2011), gastric carcinoma (Cheng et al., 2010) and lung adenocarcinoma. (Lv et al., 2015). Viceversa, in colorectal cancer, overexpression of MMP-12 is associated with increased survival of the patient, presumably as a result of an inhibitory effect on angiogenesis (Nocito et al., 2008). These data suggest that MMP-12 has a tumor suppressor role in this context.

Why is it important to know if a gene has an oncogenic role or a tumor suppressor role in a specific tumor? A careful evaluation of the functional status of a dual-role gene and the context-dependent role in response to specific treatments in different tumors might be fundamental to guide therapeutic decisions. The identification of genes with a dual role can also be used to define common genetic profiles shared by subgroups of patients who may benefit from targeted treatment strategies. For example, ERBB2 is amplified and h changing amino acid sequences. Furthermore, tumor mutation rates can be of several orders of magnitude, but it does not necessarily mean that all of the abnormal genes are also involved in the development of cancers (Lawrence et al., 2013).

Therefore, mutations and genes must be characterized together to identify biologically relevant mutations; an integration with functional enrichment analysis is essential in the interpretation of the disease.

However, to our knowledge, no study focuses on the methods to detect genes with dual role i.e. the moonlight genes. With the term ‘moonlight gene’ we mean a gene that overexpressed in 20 % to 25% of breast cancer and is the target of the monoclonal antibody trastuzumab, a drug that is effective only when ERBB2 is amplified and overexpressed (Fackenthal and Olopade, 2007).

We here propose MoonlightR as a new approach to define TSGs and OCGs based on functional enrichment analysis, inferred gene regulatory networks and upstream regulator analysis to score the importance of key biological processes that are related to cancer, such as apoptosis or cellular proliferation, with respect to the studied cancer. In addition, the Functional Enrichment Analysis (FEA) and Upstream regulator Analysis (URA), used in the Moonlight protocol, can suggest enriched biological functions for experimental validation extracting the relevant signaling pathways from differentially expressed genes.

## METHOD DETAILS

The Moonlight protocol aims:

1. To identify biological processes enriched by a gene set of differentially expressed genes (DEGs) between two conditions by means of FEA;
2. To Identify the upstream regulators of those biological processes that are enriched significantly in the comparison by means of URA;
3. To identify ‘driver genes’ with dual role that acts as TSG or OCG in different cancer contexts (i.e., different cancer types, subtypes or stages).

### Availability

The MoonlightR package is released under GPLv3 License.

MoonlightR is freely available within the Bioconductor project at http://bioconductor.org/packages/MoonlightR/.

### Data preparation and analysis

The legacy level-3 data of the PanCancer studies (18 cancer types) from the The Cancer Genome Atlas (TCGA) cohort deposited in the Genomic Data Commons (GDC) Data Portal were used in this study and downloaded in May 2017.

RNA-seq raw counts of 7364 cases (6652 TP and 712NT samples) as legacy archive, and using the reference of the hg19 genome were downloaded, normalized and filtered using the R/Bioconductor package TCGAbiolinks version 2.5.9 (Colaprico et al., 2016) using GDCquery(), GDCdownload() and GDCprepare() functions for tumor types (level 3, and platform “IlluminaHiSeq_RNASeqV2”), as well as using data.type as “Gene expression quantification” and file.type as “results”. This allowed to pull out the raw expression signal for expression of a gene for each case following the TCGA pipeline used to create Level 3 expression data from RNA Sequence data that uses MapSplice (Wang et al., 2010) to do the alignment and RSEM to perform the quantification (Li et al., 2010). Integrative analysis using mutation, clinical and gene expression were performed following our recent TCGA’s workflow (Silva et al., 2016).

For the case study n.2 and n.3 we used Breast invasive carcinoma (BRCA) from TCGA as deposited in the GDC Data Portal. In particular, RNA-seq raw counts of 1211 BRCA cases as legacy archive, and using the reference of hg19 were downloaded, normalized and filtered using the R/Bioconductor package TCGAbiolinks following the above pipeline. Among BRCA samples 1097 were Primary solid Tumor (TP) and 114 solid Tissue Normal (NT). The aggregation of the two matrices (tumor and normal) for both tumor types was then normalized using within-lane normalization to adjust for GC-content effect on read counts and upper-quantile between-lane normalization for distributional differences between lanes applying the TCGAanalyze_Normalization() function adopting the EDASeq protocol (Risso et al., 2011) and (Bullard et al., 2010).

Molecular subtypes, mutation data and clinical data were pulled using TCGAbiolinks and the following functions: TCGAquery_subtype(), GDCquery_maf() retrieving somatic variants that were called by the MuTect2 pipeline, and GDCquery_clinic() respectively.

For BRCA tumors with PAM50 classification (Ciriello et al., 2015) were stratified in 5 molecular subtypes: Basal-like (98), HER2-enriched (58), Luminal A (231), Luminal B (127) and Normal-like (8). Normal-like samples were not considered in this analysis due for the limited number of samples availability.

Tumor stages information was retrieved from the clinical data grouping to main stages (I,II,III,IV) each subgroups Ia, IIb, IIIc etc.

### Databases

Gene Programs used for Moonlight analysis were expert-based selected considering biological functions enriched by the PanCancer comparison, using the differential expressed genes between tumor samples and normal samples.

Furthermore we used the Molecular Signatures Database (MSigDB) considering the following collections to perform the Gene Set Enrichment Analysis (GSEA) analysis. (Subramanian et al., 2005).

| Collections | FileName |
| --- | --- |
| H: hallmark gene sets | h.all.v6.1.symbols.gmt.txt |
| C2: BIOCARTA pathway database | c2.cp.biocarta.v6.1.symbols.gmt.txt |
| C2: KEGG pathway database | c2.cp.kegg.v6.1.symbols.gmt.txt |
| C2: REACTOME pathway database | c2.cp.reactome.v6.1.symbols.gmt.txt |
| C3 TFT: transcription factor targets | c4.all.v6.1.symbols.gmt.txt |
| C5 BP: GO biological process | c5.bp.v6.1.symbols.gmt.txt |
| C5 CC: GO cellular component | c5.cc.v6.1.symbols.gmt.txt |
| C5 MF: GO molecular function | c5.mf.v6.1.symbols.gmt.txt |
| C6: oncogenic signatures | c6.all.v6.1.symbols.gmt.txt |
| C7: immunologic signatures | c7.all.v6.1.symbols.gmt.txt |

### Gold-standard of known driver genes (OCG and TSG)

A recent review has argued that a comparative assessment of role prediction methods is not straightforward due to the lack of a clear gold-standard of known OCGs and TSGs (Tokheim et al., 2016). To create the best currently available training set of known OCGs and TSGs we added those genes to our training set that are verified by at least two sources.

We retrieved a list of validated OCGs and TSGs from the Catalogue of somatic mutations in cancer (COSMIC) (Forbes et al., 2008). The list consists of 84 OCGs, 55 TSGs, 17 dual role genes and 439 genes without validated role. The list provides additional information such as the type of mutation dominant (448), recessive (134), dominant/recessive (7) and undeclared (3). Crossing these two types of information shows that all OCGs are linked to dominant mutation and most TSGs are linked to recessive mutations (42), nine are linked to dominant mutations.

A second list was published in (Vogelstein et al., 2013), it consists of 54 OCGs and 71 TSG.

### Functionalities overview

The Moonlight pipeline tool / suite provides well-known and new functionalities as illustrated in (Figure 1), in particular here follows the details and each of the functions to be employed:

1. **Data Collection using the functions getDataTCGA** & **getDataGEO**, allow the end-user to retrieve gene expression data from TCGA / GDC and GEO data portal, respectively.
2. **D** ifferential **P**henotype **A**nalysis (**DPA**) to identify genes or regions that are significantly different when two conditions are compared such as normal and tumor, or normal and a certain cancer stage, or even normal and a specific molecular cancer subtype.
3. **F** unctional **E**nrichment **A**nalysis (**FEA**), using Fisher’s test, to identify gene sets (with biological functions linked to the cancer study) and that are significantly enriched in the regulated genes (RGs). The principle behind FEA consists in i) evaluate if some of those DEGs are involved in a BP through an assessment of the overlap between the list of DEGs and genes relevant to this BP determined by literature mining; ii) detect the BPs mainly enriched by DEGs a Fisher Exact Test is used to calculate the probability of each BP is enriched by the entire list of DEGs.
4. **G**ene **R**egulatory **N**etwork (**GRN**) inferred between each single DEG (sDEG) and all genes by means of mutual information, obtaining for each DEG a list of regulated genes (RG).
5. **U**pstream **R**egulator **A**nalysis (**URA**) for DEGs. In each enriched gene set, we defined a Moonlight Z-score as the ratio between the sum of all predicted effects for all the gene involved in the specific function and the square-root of the number of all genes.
6. **P**attern **R**ecognition **A**nalysis (**PRA**) identifies candidate TSGs (down) and OCGs (up). We either use user defined biological processes or random forests.
7. We applied the above procedure to multiple cancer types to obtain cancer-specific lists of TCGs and OCGs. We compared the lists for each cancer: if a sDEG was TSG in a cancer and OCG in another we defined it as dual-role TSG-OCG. Otherwise if we found a sDEG defined as OCG or TSG only in one tissue we defined it tissue specific biomarker.
8. We used the COSMIC database to define a list of gold standard TSG and OCGs to assess the accuracy of the proposed methods.

**Figure 1.**
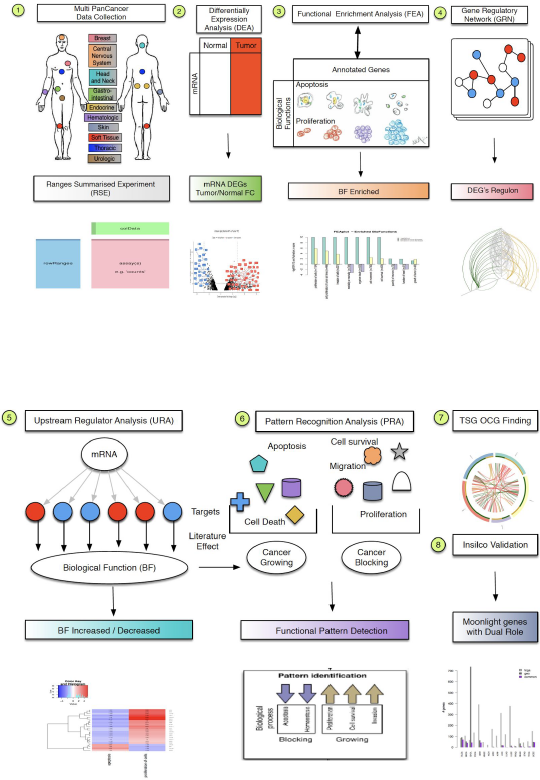
Moonlight pipeline. Proposed Moonlight functionalities for functional biological process and driver genes discovery.

### ROMA score for pathway activity

For the pathway activity evaluation, Representation and quantification Of Module Activity (ROMA) (See https://github.com/sysbio-curie/Roma) (Martignetti et al., 2016) was also employed as alternative to the Moonlight Z-score. For each module under analysis, the algorithm applies Principal Component Analysis (PCA) to the sub-matrix composed of the expression values of the signature genes across samples. Then ROMA evaluates the module overdispersion by verifying if the amount of variance explained by the first principal component of the expression sub-matrix (L1 value in ROMA) is significantly larger than that of a random set of genes of the same size. This thus represents an unsupervised approach, that can be also used in combination to the supervised Z-score to detect concordant signals.

## RESULTS

### Case study n.1a (FEA using TCGA BRCA gene expression)

In this case study using we defined a gene expression signature (GeS) after running TCGAanalyzeDEA between breast cancer samples BRCA TP and NT obtained 3390 DEGs namely the differentially expressed genes (DEGs). Figure 2 showed the top 32 BPs enriched significantly with |Moonlight-score| >=1 and FDR <= 0.01 using the above GeS. Increased levels reported in yellow and decreased in purple, and in green showing the -logFDR / 10. A negative Z-score indicates that the process’ activity is decreased. A positive Z-score indicates that the process’ activity is increased.

**Figure 2.**
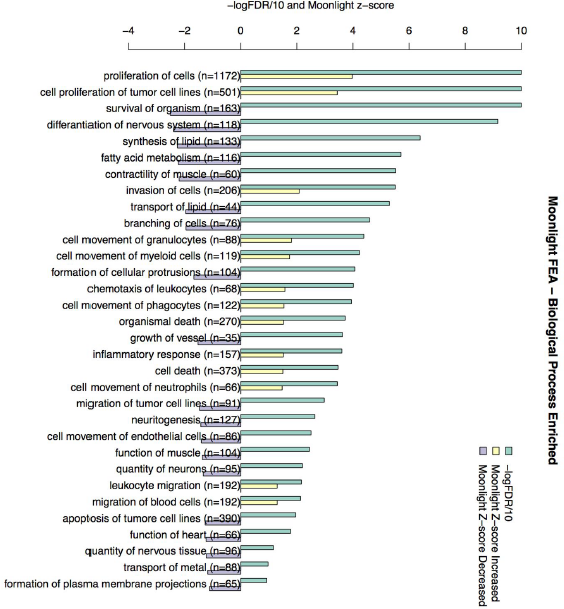
Case study n.1a (Functional Enrichment Analyisis) using TCGA BRCA gene expression. Barplot showing the top 32 BPs enriched significantly with |Moonlight-score| >=1 and FDR <= 0.01. Increased levels reported in yellow and decreased in purple, and in green showing the -logFDR / 10. A negative Z-score indicates that the process’ activity is decreased. A positive Z-score indicates that the process’ activity is increased.

### Case study n.1b (Upstream regulator analysis (URA) using TCGA BRCA gene expression)

We performed Upstream regulator analysis (URA) using the GeS (n=3390 DEGs) defined in case study n.1a to identify the cause of a downstream effect in cellular biology. The identification of upstream regulator molecules and their related biological processes can help to better understand the detected expression changes by the DEA.

In the figure below (Figure 3) we applied also the Pattern recognition Analysis on the URA obtained from the BRCA considering as apoptosis of tumor cell lines and proliferation of cells as the interested biological process for the definition of upstream regulators.

**Figure 3.**
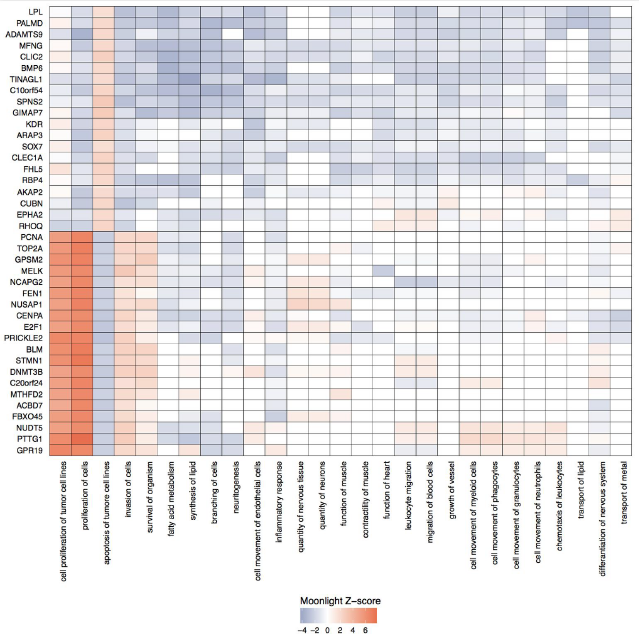
Case study n.1b (Upstream regulator analysis (URA) using TCGA BRCA gene expression). Heatmap showing to 40 differentially expressed upstream regulators of the biological process from case study n.1a. applying Hierarchical cluster analysis based after computing the distance matrix by euclidean method.

### Case study n.1c (Quantification of genes effect on Biological Functions (BF) data-driven by ROMA-score

We computed the activity score of biological processes potentially modulated by OCG and TSG using ROMA (Representation and quantification Of Module Activities) (Martignetti et al., 2016), a gene-set-based quantification algorithm, for more details see Methods. As a case study, we applied ROMA to the BRCA dataset of TCGA (table: module_scores.xls). This approach revealed substantial similarity with the results obtained by Moonlight Z-score.

### Case study n. 2: Characterize intertumoral (between tumors) genomic epigenomic and transcriptomic heterogeneity in cancer tissue by identifying dual role genes

In this case study, we used the expression levels of genes in all the samples obtained from TCGA / GDC with IlluminaHiSeq RNASeqV2 in 18 normal tissues (NT) and 18 cancer tissues (CT) according to the TCGA criteria (Table 1). We applied the complete Moonlight pipeline (FEA-URA-PRA) to extract those genes that were significantly increasing or decreasing biological processes such as proliferation of cells and apoptosis.

**Table 1.** TCGA and GEO PanCancer data used in this study.

| TCGA |  |  |  | GEO |  |  |  |  | Validation |
| --- | --- | --- | --- | --- | --- | --- | --- | --- | --- |
| Cancer | TP | NT | DEG* | Dataset | TP | NT | Platform | DEG** | Common |
| BLCA | 408 | 19 | 2937 | GSE13507 | 165 | 10 | GPL65000 | 2099 | 896 |
| BRCA | 1097 | 114 | 3390 | GSE39004 | 61 | 47 | GPL6244 | 2449 | 1248 |
| CHOL | 36 | 9 | 5015 | GSE26566 | 104 | 59 | GPL6104 | 3983 | 2587 |
| COAD | 286 | 41 | 3788 | GSE41657 | 25 | 12 | GPL6480 | 3523 | 1367 |
| ESCA | 184 | 11 | 2525 | GSE20347 | 17 | 17 | GPL571 | 1316 | 406 |
| GBM | 156 | 5 | 4828 | GSE50161 | 34 | 13 | GPL570 | 4504 | 2660 |
| HNSC | 520 | 44 | 2973 | GSE6631 | 22 | 22 | GPL8300 | 142 | 129 |
| KICH | 66 | 25 | 4355 | GSE15641 | 6 | 23 | GPL96 | 1789 | 680 |
| KIRC | 533 | 72 | 3618 | GSE15641 | 32 | 23 | GPL96 | 2911 | 939 |
| KIRP | 290 | 32 | 3748 | GSE15641 | 11 | 23 | GPL96 | 2020 | 756 |
| LIHC | 371 | 50 | 3043 | GSE45267 | 46 | 41 | GPL570 | 1583 | 860 |
| LUAD | 515 | 59 | 3498 | GSE10072 | 58 | 49 | GPL96 | 666 | 555 |
| LUSC | 503 | 51 | 4984 | GSE33479 | 14 | 27 | GPL6480 | 3729 | 1706 |
| PRAD | 497 | 52 | 1860 | GSE6919 | 81 | 90 | GPL8300 | 246 | 149 |
| READ | 94 | 10 | 3628 | GSE20842 | 65 | 65 | GPL4133 | 2172 | 1261 |
| STAD | 415 | 35 | 2622 | GSE2685 | 10 | 10 | GPL80 | 487 | 164 |
| THCA | 505 | 59 | 1994 | GSE33630 | 60 | 45 | GPL570 | 1451 | 781 |
| UCEC | 176 | 24 | 4183 | GSE17025 | 11 | 53 | GPL570 | 2487 | 955 |

(Figure 4) showed the results of the moonlight pipeline for case study n.2 that can be visualized with a circos plot using the function plotCircos.

**Figure 4.**
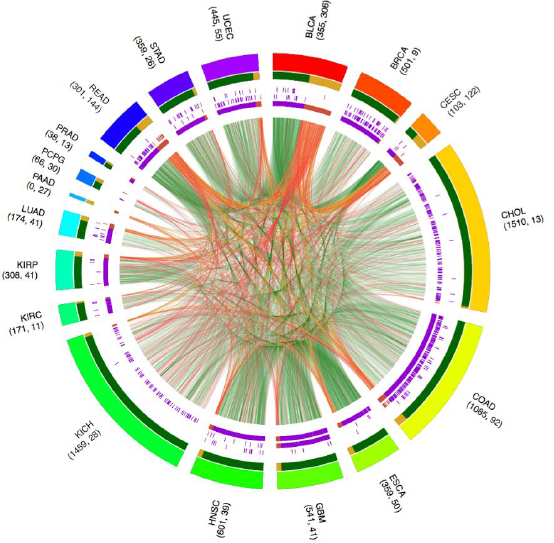
Case study n. 2: Characterize intertumoral (between tumors) genomic epigenomic and transcriptomic heterogeneity in cancer tissue by identifying dual role genes. Circos Plot showing an integrative analysis of 18 TCGA cancer types. Outer ring: color by cancer type, Inner ring: OCGs and TSGs, Inner connections: green: common OCGs yellow: common TSGs red: possible dual role. Reporting in purple, orange, the number for each stage, mutations inframe deletion, inframe insertion, missense.

### Case study n.3: Characterize intratumoral (within a tumor) genomic epigenomic and transcriptomic heterogeneity in cancer tissue by identifying dual role genes

In this case study Figure 5 (a) Comparison between BRCA normal samples and specific stages samples. (‘c) Comparison between normal samples and specific BRCA molecular subtypes samples. (b) (d) Circos plots for stages and molecular subtypes respectively, of moonlight genes predicted using expert knowledge PRA with two selected biological processes (BP) such as apoptosis and proliferation of cells. Outer to inner layer OCG in green TSG in yellow, reporting in purple, orange, the number for each stage, mutations inframe deletion, inframe insertion, missense. Connecting gene-gene links between two cancer stages, green in OCG in both, yellow TSG in both, red moonlight genes playing dual role OCG in one and TSG in the other.e data download.

**Figure 5:**
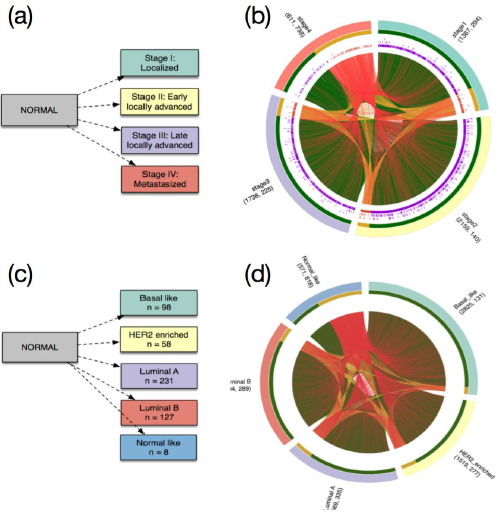
Case study n.3: Characterize intratumoral (within a tumor) genomic epigenomic and transcriptomic heterogeneity in cancer tissue by identifying dual role genes. (a) Comparison between BRCA normal samples and specific stages samples. (‘c) Comparison between normal samples and specific BRCA molecular subtypes samples. (b) (d) Circos plots for stages and molecular subtypes respectively, of moonlight genes predicted using expert knowledge PRA with two selected biological processes (BP) such as apoptosis and proliferation of cells. Outer to inner layer OCG in green TSG in yellow, reporting in purple, orange, the number for each stage, mutations inframe deletion, inframe insertion, missense. Connecting gene-gene links between two cancer stages, green in OCG in both, yellow TSG in both, red moonlight genes playing dual role OCG in one and TSG in the other.

## Conclusion

In this paper, we proposed a new methodology called Moonlight to identify driver cancer genes and their specific role in different contexts. In particular, our method can predict oncogenes (or tumor suppressor genes) as those genes that increase (decrease) biological processes related to cancer growth while at the same time decrease (increase) biological processes that are blocking cancer growth. Moreover, our approach sounds promising to identify those genes with a dual role i.e., the moonlight genes. These genes are emerging in cancer studies and they are elusive to identify due to their context-dependent role, i.e. they could act as an oncogene in a certain cancer type or stage, as well as a tumor suppressor in other contexts. They thus challenges the development of targeted therapies and treatments, emphasizing the needs of new methods to accurately identifying them. Examples of processes related to cancer growth included in our study are cell proliferation and cell survival, examples of processes related to cancer blocking are homeostasis and cell death. A process is increased (decreased) if the associated functional enrichment analysis (FEA) yields positive (negative) Z-score values, i.e. high correlation (high anti-correlation) between the gene expression pattern and the literature-curated information. Then we determine whether a gene is increasing (decreasing) the biological process using an inferred gene regulatory network and subsequent Upstream Regulator Analysis (URA)

## Acknowledgements

We are grateful to Matthieu Defrance, Kridsadakorn Chaichoompu, Kristel Van Steen, Benjamin Haibe-Kains for suggestions and scientific advices in the Moonlight project. The project was supported by the BridgeIRIS project [http://mlg.ulb.ac.be/BridgeIRIS], funded by INNOVIRIS, Region de Bruxelles Capitale, Brussels, Belgium, and by GENGISCAN: GENomic profiling of Gastrointestinal Inflammatory-Sensitive CANcers, [http://mlg.ulb.ac.be/GENGISCAN] Belgian FNRS PDR (T100914F to A.C., C.O. and G.B.) EP group was supported by a grant from LEO Foundation (grant number LF17006) and InnovationFund Denmark (grant number 5189-00052B).

